# Improved protein contact prediction using dimensional hybrid residual networks and singularity enhanced loss function

**DOI:** 10.1101/2021.05.10.443415

**Authors:** Yunda Si, Chengfei Yan

## Abstract

Deep residual learning has shown great success in protein contact prediction. In this study, a new deep residual learning-based protein contact prediction model was developed. Comparing with previous models, a new type of residual block hybridizing 1D and 2D convolutions was designed to increase the effective receptive field of the residual network, and a new loss function emphasizing the easily misclassified residue pairs was proposed to enhance the model training. The developed protein contact prediction model referred to as DRN-1D2D was first evaluated on 105 CASP11 targets, 76 CAMEO hard targets and 398 membrane proteins together with two in house-developed reference models based on either the standard 2D residual block or the traditional BCE loss function, from which we confirmed that both the dimensional hybrid residual block and the singularity enhanced loss function can be employed to improve the model performance for protein contact prediction. DRN-1D2D was further evaluated on 39 CASP13 and CASP14 free modeling targets together with the two reference models and six state-of-the-art protein contact prediction models including DeepCov, DeepCon, DeepConPred2, SPOT-Contact, RaptorX-Contact and TripleRes. The result shows that DRN-1D2D consistently achieved the best performance among all these models.

## 1 Introduction

The knowledge of protein residue-residue contacts is valuable for predicting protein structures with no-homologous templates, therefore, protein contact prediction had become a long-standing scientific problem for decades [1]. In recent years, great progress has been made on solving this problem due to the development of coevolutionary analysis methods and deep learning methods to predict protein contacts [2,3]. Coevolutionary analysis methods generally predict protein contacts through inferring residue pairs with direct evolutionary couplings from the multiple sequence alignment (MSA) of homologous proteins [4–8]. However, coevolutionary analysis methods can only accurately predict protein contacts when a large number of homologous sequences of the protein are available to produce an accurate estimation of the unknown parameters of the statistical models used in coevolutionary analysis. Deep learning methods apply deep learning models trained from large datasets of proteins to predict protein contacts, in which the coevolution and other properties of the protein obtained from the MSA of homologous proteins are often used as the input features. Deep learning methods are less dependent on the number of homologous sequences because parameters of the deep learning models are predetermined through minimizing the loss between the predicted protein contact maps and the experimental protein contact maps in large protein datasets.

Many deep learning models based on various neural network architectures including long short-term memory networks (LSTM) [9], deep belief networks (DBN) [10,11], generative adversarial networks (GAN) [12], convolutional neural networks (CNN) [13], residual neural networks (ResNet) [14], etc. have been recently developed. Among these developed models, the deep residual learning models showed great success, since most of the top protein contact prediction models in recent Critical Assessment of protein Structure Prediction (CASP) employed or partially employed residual neural networks [2]. Although some models based on other network architectures can also achieve comparable performance. The first deep residual learning-based protein contact prediction model was developed by Xu and his coworkers, in which two residual networks were successively applied to transform the input features for protein contact prediction [14]. The first is a 1D residual network built by 1D residual blocks to conduct 1D convolutional transformations of the 1D sequential input features; the second is a 2D residual network built by 2D residual blocks to conduct 2D convolutional transformations of the 2D pairwise features and the 2D transformed feature maps of the 1D sequential feature maps output from the 1D residual network. After Xu’s work, deep residual learning was also applied by other groups to develop protein contact prediction models, in which efforts on improving the model performance include using much deeper residual networks, increasing the size of the training dataset, engineering new input features, employing metagenome sequence data and etc [15–19]. Most of these models only employed 2D residual blocks to build the residual network, in which the 2D transformed 1D sequential features were directly combined with the 2D pairwise features to form the input features of the residual network, or only the 2D pairwise features were used as the input features considering the pairwise features are generally much more important than the sequential features. It is clear that residual networks built by 2D residual blocks are effective network architectures for protein contact prediction, but whether the model performance can be further improved by designing novel residual blocks is unknown. Besides, appropriately calculating the training loss is also important for the model training. Most previous works applied the binary cross entropy (BCE) loss function to calculate the training loss, in which residue pairs in the protein contact map are equally weighted in the loss calculation. However, residue pairs in the contact-sparser regions of the protein contact map are more difficult to be correctly classified, emphasizing the easily misclassified residue pairs may enhance the model training.

In this study, we present a new deep residual learning-based protein contact prediction model. Comparing with previous models, a new type of residual block hybridizing 1D and 2D convolutions was designed to build the residual network, with which we show that the effective receptive field of the residual network can be significantly increased. Besides, a new loss function emphasizing the easily misclassified residue pairs was proposed to enhance the model training. The model was trained on the Xu’s original training dataset [14] for developing the first deep residual learning-based protein contact model, which is referred to as DRN-1D2D in this study. Besides, we also developed two additional models on the same dataset as the reference, in which one model was based on the basic 2D residual block and the BCE loss function, and another was based on the dimensional hybrid residual block and the BCE loss function. We first evaluated DRN-1D2D and the two reference models on Xu’s original three test sets (105 CASP11 proteins, 76 CAMEO hard targets and 396 membrane proteins) [14]. The result shows that DRN-1D2D consistently outperformed the two reference models, and between the two reference models, the model based on the new residual block consistently outperformed the model based on the basic 2D residual block, which confirms that the dimensional hybrid residual block and the singularity enhanced loss function can both be employed to improve the model performance for protein contact prediction. Besides, DRN-1D2D also consistently outperformed Xu’ original model, although these two models were developed on the same dataset and also have the same input features. We further evaluated DRN-1D2D on 39 CASP13 and CASP14 free modeling targets together with the two in house-developed reference models and six state-of-the-art protein contact prediction models including DeepCov [15], DeepCon [18], DeepConPred2 [20], SPOT-contact [9], RaptorX-Contact [21] and TripleRes [22]. The result illustrates that DRN-1D2D consistently achieved the best performance among these models. It is worth noting that comparing with DRN-1D2D, both RaptorX-Contact and TripleRes have much larger network sizes and were also trained on much larger protein datasets.

## 2 Materials and Methods

### 2.1 The network architecture of DRN-1D2D

#### 2.1.1 The input feature maps

The input features of DRN-1D2D include three 1D sequential features: the position-specific scoring matrix (PSSM), the predicted 3-state secondary structure matrix (SS3), and the predicted 3-state solvent accessibility matrix (ACC), and four 2D pairwise features: the evolutionary coupling matrix from CCMpred [23], the mutual information matrix, the APC-corrected mutual information matrix and the pairwise contact potentials matrix, which are the same with Xu’s original model [14]. The 1D sequential features are first concatenated along the columns. Let V = {*v*_1_, *v*_2_,…, *v*_*i*_,…, *v*_*L*_} be the concatenated 1D sequential feature matrix where L is the sequence length and *v*_*i*_ is a vector storing the 1D features for residue i. A feature vector for residue pair ij can be created by concatenating 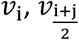 and *v*_j_ to a single vector, with which we can convert the 1D sequential feature matrix to 2D pairwise feature maps. The 2D converted feature maps (L*L*78) of the 1D sequential features and the 2D pairwise feature maps (L*L*4) are further concatenated to form the input feature maps (L*L*82) of the network of DRN-1D2D.

#### 2.1.2 The residual network

The network architecture of DRN-1D2D is shown in Figure 1A. First, a 1*1 convolutional layer followed by an instance normalization (IN) layer and a LeakyReLU activation layer is used to transform the number of channels of the input feature maps from 82 to 64. Then 25 residual blocks are used to transform the 64 channels of feature maps. Finally, another 1*1 convolutional layer is applied to transform the number of channels from 64 to 1, which is followed by the sigmoid transformation to produce the predicted contact map. In this study, instead of the 2D residual blocks, a new type of residual block hybridizing 1D and 2D convolutions was designed to build the residual network, which is described in the following section.

#### 2.1.3 The dimensional hybrid residual block

Receptive field is defined as the spatial extent of the inputs used in the calculation of an output unit, and the input region outside the receptive field of the output unit does not affect the output value of that unit. The receptive field size is a crucial issue for designing convolutional neural networks, for the receptive field of the network needs to be large enough to capture the complex information in the input feature maps. Luo et al. show that the effective area in the receptive field only occupies a fraction of the theoretical receptive field, for the distribution of impact in the receptive field of 2D convolutional neural networks distributes as a gaussian, which generally decays quickly from the center [24]. In this study, to increase the effective receptive field of the residual network, two additional 1D convolutional branches with kernel sizes of 9*1 and 1*9 are added to the basic 2D residual block formed by 3*3 convolution kernels (see Figure 1B and 1C, referred to as 2D block and 1D2D block in this study). Specifically, given an input of the residual block (*x*_*i*_), the input is first transformed successively by the first convolution followed by instance normalization and LeakyReLU activation, and the second convolution followed by instance normalization from each branch separately; then the outputs from the three branches are summed together and added to the input (*x*_*i*_ + *f*_1*9_(*x*_*i*_) + *f*_3*3_(*x*_*i*_) + *f*_9*1_(*x*_*i*_)), which is finally transformed by the LeakyReLU activation to produce the output of the residual block. The dimensions of the input feature maps are kept in the convolutions through the application of zero paddings symmetrically on each feature map.

In Figure 2, we show the distribution of impact in the receptive field of the residual network built by 25 new residual blocks (i.e. the network of DRN-1D2D), by 25 basic 2D residual blocks, by 75 basic 2D residual blocks and by 142 basic 2D residual blocks respectively. The impact of each input pixel in the receptive field is evaluated by its input gradient when the convolution kernels are constant kernels. As we can see from Figure 2 that the residual network built by 25 new residual blocks has a much larger effective receptive field than the residual network built by 25 and 75 basic 2D residual blocks, and has a similar effective receptive field to the residual network built by 142 basic 2D residual blocks. The effective receptive field of the residual network built by 75 basic 2D residual blocks is shown as a reference for which has the same number of parameters with the network of DRN-1D2D (see Table S1).

### 2.2 The model training of DRN-1D2D

#### 2.2.1 The calculation of training loss

Protein contact maps are sparse matrices in which contacting residue pairs are labeled with ones and non-contacting residue pairs are labeled with zeros. A pair of residues are generally considered to be in contact if their *C*_*β*_ -*C*_*β*_ distance (*C*_*α*_ -*C*_*α*_ distance for GLY) is smaller than 8.0Å. A protein contact map generally contains much more zeros than ones, and regions that further away from the diagonal of the contact map is often sparser, for the residues separated by more residues have lower contact probabilities. Training a deep learning model for protein contact prediction is the process to determine the parameters of the network through minimizing the loss between the predicted protein contact maps and the real protein contact maps in the training set. In practice, due to the memory limitation, the parameters are usually updated successively based on the loss of each protein contact map (i.e. batch size equals to 1) across the training set, which is often calculated with the BCE loss function:

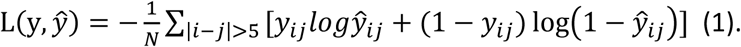

Where y represents the true contact map; *ŷ* represents the prediction; N is the total number of the non-local residue pairs (residues separated by more than five residues). The local residue pairs are often not included in the loss calculation for the local residue contacts are trivially formed due to the peptide chain connection. In the BCE loss function, residue pairs in different regions of contact map are equally weighted. However, residue pairs in the contact-sparser regions (e.g. long-range or extra long-range contacts) of the protein contact map are more difficult to be correctly classified, therefore which should be emphasized in the model training. In this study, we propose a singularity enhanced loss function to increase the weight of these residue pairs:

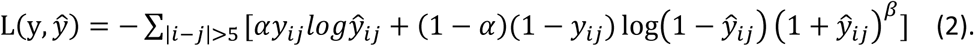

Comparing with the BCE loss, first, a parameter *α* is introduced to weight the contacting and non-contacting residue pairs; second, (1 + *ŷ*_*ij*_)^*β*^ is introduced to increase the loss contribution from the misclassified non-contacting residue pairs. Since residue pairs in the contact-sparser regions of the contact map are more difficult to be correctly classified, the introduction of this term can automatically emphasize these residue pairs. Besides, in the BCE loss, the mean of the losses across all non-local residue pairs are used in calculating the loss of each contact map, but in our proposed loss function, the sum rather than the mean is used to calculate the loss of each contact map, for we would like larger proteins to play a more important role in the model training. The two parameters *α* and *β* were carefully calibrated on a relative smaller dataset, and we found that when 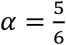 and *β*=3, the obtained model achieved the best performance, therefore which were used in this study for the development of DRN-1D2D.

#### 2.2.2 The training protocol

We directly trained the model of DRN-1D2D on the dataset of 6767 proteins (including all the input features and protein contact maps) prepared by Xu for developing the first deep residual learning-based protein contact prediction model [14]. Similar to Xu’s protocol, seven different models were trained separately through seven-fold cross-validation from the dataset, and the final model is an average of the seven models. The He initialization [25] was used to initialize the parameters of the network, and the parameters were optimized by the AdamW optimizer with a weight decay of 0.1 and an initial learning rate of 0.001. In each epoch of training, the network parameters were successive updated based on the loss of each contact map across the training set calculated by the singularity enhanced loss function (i.e. batch size=1). If the total loss did not decrease within 2 epochs, the learning rate was decayed to 0.1 of its original value. After the learning rate decayed twice, the training was stopped, and the model with the lowest total loss from the previous epoch was saved as the final model for protein contact prediction (see Figure S1). The model was built with PyTorch and trained on the NVIDIA Tesla P100. Due to the GPU memory limitation, for the protein with sequence length larger than 400, a protein fragment with length equaling to 400 randomly selected from the protein was used in the model training. The average time for training a model is about 1 day. Besides, we also trained two additional models using the above protocol on the same dataset except that: in one model the residual blocks in the residual network were replaced by the basic 2D residual blocks and the training loss was calculated by the BCE loss function; in another model, the dimensional hybrid residual network was kept but still the model was trained with the BCE loss. These two models referred to as BCE-2D and BCE-1D2D were used as the reference models in this study.

### 2.3 The model evaluation of DRN-1D2D

Xu’s original three test sets (105 CASP11 targets, 76 CAMEO targets and 398 membrane proteins) and the free modeling targets of CASP13 and CASP14 with released native structures (24 targets from CASP13 and 15 targets from CASP14) were used respectively to evaluate the performance of DRN-1D2D. For Xu’s original test sets, the input features prepared by Xu were directly used by DRN-1D2D for protein contact prediction, and the two additional in house-developed models (BCE-2D and BCE-1D2D) employing the same set of input features were used as the references.

For the CASP13 and CASP14 datasets, DeepMSA was applied to successively search Uniclust30 (2018-08), Uniref90 (2019-11) and Metaclust (2018-06) to build the MSA of homologous proteins for each target, and the input features of DRN-1D2D were generated from the MSA according to the protocols described by Xu (https://github.com/j3xugit/RaptorX-Contact). A detailed description of the preparation of the input features for DRN-1D2D was also provided in the supplementary Text S1. Besides, the two in house-developed reference models (BCE-2D and BCE-1D2D) and six state-of-the-art outside protein contact prediction models including DeepCov (https://github.com/psipred/DeepCov), DeepCon (https://github.com/ba-lab/DEEPCON), DeepConPred2 (https://github.com/THU-gonglab/DeepConPred2), SPOT-contact (https://sparks-lab.org/server/spot-contact/), RaptorX-Contact (https://github.com/j3xugit/RaptorX-Contact) and TripleRes (https://zhanglab.ccmb.med.umich.edu/TripletRes/) were also evaluated on CASP datasets for the purpose of comparison, in which the two in house-developed reference models employed the same set of input features as DRN-1D2D, and the six state-of-the-art outside models employed the same set of MSAs as DRN-1D2D to produce their input features.

## 3. Results and Discussion

### 3.1 The performance of DRN-1D2D on Xu’s original three test sets

We first evaluated DRN-1D2D and the two reference models (BCE-2D and BCE-1D2D) on Xu’s original three test sets (105 CASP11 targets, 76 CAMEO targets and 398 membrane proteins), and the input features prepared by Xu were directly used by DRN-1D2D and the two reference models for protein contact prediction. In order to use the performance of Xu’s original model as an extra reference, we used the same evaluation method as Xu to evaluate the performances of our models. Specifically, the accuracies of the top L/K (k=10, 5, 2, 1) of the short-, medium- and long-range predicted contacts were used to evaluate the performance of each model, where L is the protein sequence length [14]. A pair of residues are considered to be in contact if their *C*_*β*_-*C*_*β*_ distance (*C*_*α*_-*C*_*α*_ distance for GLY) is smaller than 8.0Å, and a contact is defined to be short-, medium- and long-range if the sequence distance of the two residues falls into [6, 11], [12, 23] and ≥ 24. In Table 1, we show the mean precisions of the contacts predicted by DRN-1D2D, the two reference models and Xu’s original model on the three datasets. It is worth noting that the input features for all the four models are exactly the same. As we can see from Table 1, DRN-1D2D consistently outperformed all other models, and between the two reference models, the model based on the new residual block achieved better performances than the model based on the basic 2D residual block. The accuracy comparisons between DRN-1D2D and the two reference models on the top L/2 and L/5 of the predicted long-range contacts for all individual targets from the three test sets are shown in Figure 3. As we can see from Figure 3 that the improvements actually occur consistently across most of the targets, which confirms that both the dimensional hybrid residual block and the singularity enhanced loss function can be employed to improve the model performance for protein contact prediction.

**Table 1.** The mean precisions of contacts predicted by different models on Xu’s original three test sets

| Test sets | Method | Short |  |  |  | Medium |  |  |  | Long |  |  |  |
| --- | --- | --- | --- | --- | --- | --- | --- | --- | --- | --- | --- | --- | --- |
|  |  | L/10 | L/5 | L/2 | L | L/10 | L/5 | L/2 | L | L/10 | L/5 | L/2 | L |
| CASP11 | Xu 2017 | 0.82 | 0.70 | 0.46 | 0.28 | 0.85 | 0.76 | 0.55 | 0.35 | 0.81 | 0.77 | 0.68 | 0.55 |
|  | BCE-2D | 0.83 | 0.72 | 0.47 | 0.29 | 0.83 | 0.75 | 0.55 | 0.36 | 0.79 | 0.77 | 0.66 | 0.54 |
|  | BCE-D2D | 0.84 | 0.73 | 0.48 | 0.29 | 0.84 | 0.76 | 0.56 | 0.36 | 0.83 | 0.79 | 0.69 | 0.56 |
|  | DRN-1D2D | **0.86** | **0.75** | **0.49** | **0.29** | **0.86** | **0.78** | **0.58** | **0.37** | **0.86** | **0.82** | **0.73** | **0.59** |
| CAMEO | Xu 2017 | 0.67 | 0.57 | 0.37 | 0.23 | 0.69 | 0.61 | 0.42 | 0.28 | 0.69 | 0.65 | 0.55 | 0.42 |
|  | BCE-2D | 0.69 | 0.59 | 0.41 | 0.25 | 0.68 | 0.60 | 0.44 | 0.30 | 0.67 | 0.63 | 0.53 | 0.41 |
|  | BCE-1D2D | 0.72 | 0.60 | 0.41 | 0.25 | 0.72 | 0.63 | 0.46 | 0.30 | 0.69 | 0.65 | 0.55 | 0.43 |
|  | DRN-1D2D | **0.75** | **0.63** | **0.43** | **0.26** | **0.74** | **0.65** | **0.47** | **0.31** | **0.74** | **0.69** | **0.59** | **0.46** |
| Mems | Xu 2017 | 0.60 | 0.46 | 0.27 | 0.16 | 0.66 | 0.53 | 0.33 | 0.22 | 0.78 | 0.73 | 0.62 | 0.47 |
|  | BCE-2D | 0.62 | 0.49 | 0.30 | 0.18 | 0.67 | 0.55 | 0.36 | 0.23 | 0.77 | 0.73 | 0.62 | 0.48 |
|  | BCE-1D2D | 0.64 | 0.50 | 0.31 | 0.18 | 0.67 | 0.56 | 0.36 | 0.23 | 0.78 | 0.74 | 0.64 | 0.49 |
|  | DRN-1D2D | **0.65** | **0.52** | **0.32** | **0.19** | **0.70** | **0.58** | **0.38** | **0.24** | **0.81** | **0.77** | **0.67** | **0.53** |

Note: The highest accuracy in each category is highlighted in bold font.

### 3.2 The performance of DRN-1D2D on recent CASP targets

DRN-1D2D was further evaluated on recent CASP targets. Specifically, the free modeling targets of CASP13 and CASP14 with released native structures (24 targets from CASP13 and 15 targets from CASP14) were used to evaluate the performance of DRN-1D2D. For each target, we applied DeepMSA [26] to successively search Uniclust30 (2018-08) [27], Uniref90 (2019-11) [28] and Metaclust (2018-06) [29] to build the MSA of its homologous proteins, from which we generated the input features for DRN-1D2D according to the protocols described by Xu [14]. Besides, the two in-house developed reference models and six state-of-the-art protein contact prediction models including DeepCov, DeepCon, DeepConPred2, SPOT-contact, RaptorX-Contact and TripleRes were also applied to predict the protein contacts on the same dataset. To ensure a fair comparison, for each target, the input features of all the above models were generated from the same MSA. The performance of each model was evaluated by calculating the accuracies of the top L/K (K=5, 2, 1) of the medium + long-range (sequence distance≥ 12) and long-range (sequence distance≥ 24) of the predicted contacts, which is the same with the recent CASP protocol for evaluating the performance of protein contact prediction[2]. In Table 2, we show the mean precisions of the contacts predicted by these models on the CASP13 and CASP14 targets respectively. As we can see from Table 2 that DRN-1D2D consistently achieved the best performances on both CASP13 and CASP14 targets. Besides, the accuracy comparisons between DRN-1D2D and other models on the top L/5 of the predicted medium + long-range and long-range contacts for all individual targets are shown in Figure 4 (Figure 4A-4B: DRN-1D2D versus the two in house-developed reference models; Figure 4C-4D: DRN-1D2D versus the four state-of-the-art models; for the top L/2 contact prediction, see Figure S2). As we can see from Figure 4 that DRN-1D2D consistently outperforms other models for most of the targets. It is worth noting that RaptorX-Contact is a new version of Xu’s original protein contact prediction model, which was trained on a much larger dataset containing 11410 proteins, and has a much larger network architecture; the network architecture of TripleRes is formed by four residual networks, each of the residual network has a network depth similar to DRN-1D2D [21,22].

**Table 2.**
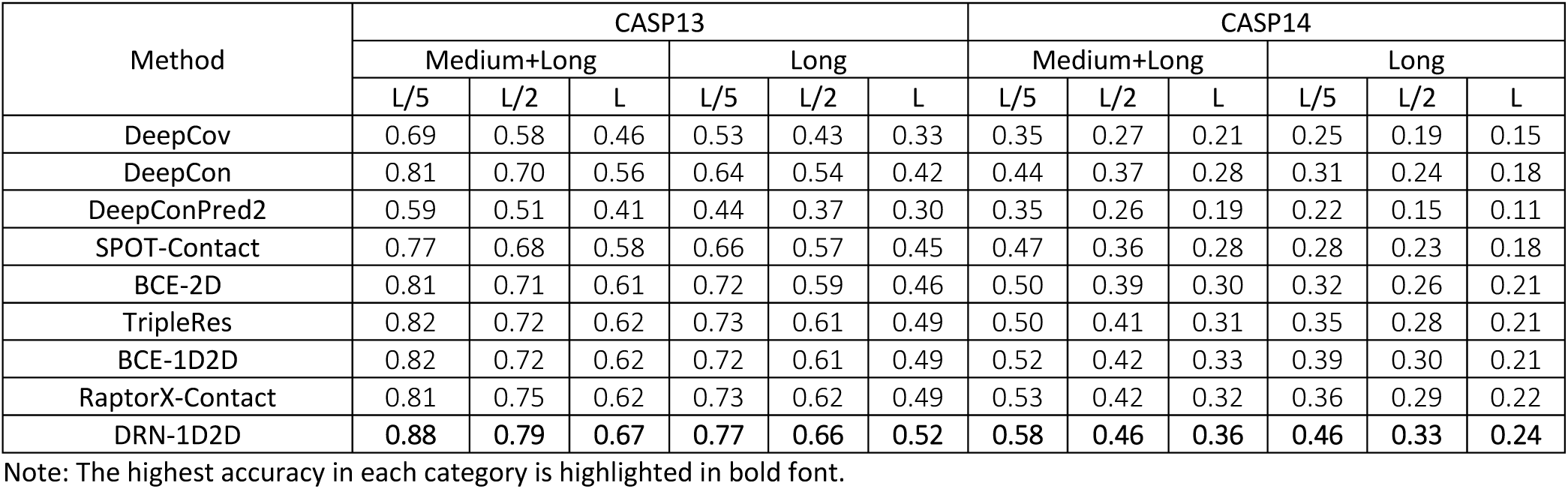
The mean precisions of contacts predicted by different models on CASP13 and14 targets

| Method | CASP13 |  |  |  |  |  | CASP14 |  |  |  |  |  |
| --- | --- | --- | --- | --- | --- | --- | --- | --- | --- | --- | --- | --- |
|  | Medium+Long |  |  | Long |  |  | Medium+Long |  |  | Long |  |  |
|  | L/5 | L/2 | L | L/5 | L/2 | L | L/5 | L/2 | L | L/5 | L/2 | L |
| DeepCov | 0.69 | 0.58 | 0.46 | 0.53 | 0.43 | 0.33 | 0.35 | 0.27 | 0.21 | 0.25 | 0.19 | 0.15 |
| DeepCon | 0.81 | 0.70 | 0.56 | 0.64 | 0.54 | 0.42 | 0.44 | 0.37 | 0.28 | 0.31 | 0.24 | 0.18 |
| DeepConPred2 | 0.59 | 0.51 | 0.41 | 0.44 | 0.37 | 0.30 | 0.35 | 0.26 | 0.19 | 0.22 | 0.15 | 0.11 |
| SPOT-Contact | 0.77 | 0.68 | 0.58 | 0.66 | 0.57 | 0.45 | 0.47 | 0.36 | 0.28 | 0.28 | 0.23 | 0.18 |
| BCE-2D | 0.81 | 0.71 | 0.61 | 0.72 | 0.59 | 0.46 | 0.50 | 0.39 | 0.30 | 0.32 | 0.26 | 0.21 |
| TripleRes | 0.82 | 0.72 | 0.62 | 0.73 | 0.61 | 0.49 | 0.50 | 0.41 | 0.31 | 0.35 | 0.28 | 0.21 |
| BCE-1D2D | 0.82 | 0.72 | 0.62 | 0.72 | 0.61 | 0.49 | 0.52 | 0.42 | 0.33 | 0.39 | 0.30 | 0.21 |
| RaptorX-Contact | 0.81 | 0.75 | 0.62 | 0.73 | 0.62 | 0.49 | 0.53 | 0.42 | 0.32 | 0.36 | 0.29 | 0.22 |
| DRN-1D2D | **0.88** | **0.79** | **0.67** | **0.77** | **0.66** | **0.52** | **0.58** | **0.46** | **0.36** | **0.46** | **0.33** | **0.24** |

486 Note: The highest accuracy in each category is highlighted in bold font.

### 3.3 The impact of MSA on the model performance of DRN-1D2D

We analyzed the impact of MSA on the model performance of DRN-1D2D. Specifically, for each CASP target, the normalized effective sequence number of the MSA 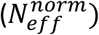 was calculated according to:

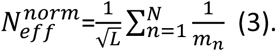

Where N is the total number of sequences in the MSA; 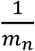 is the weight the n-th sequence with *m*_*n*_ being the number of sequences in the MSA which have a sequence identity higher than 80% to the n-th sequence; L is the sequence length. In Figure 5, we show the accuracy of the top L/5 long-range predicted contacts from DRN-1D2D versus 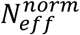 for each individual target (see Figure S3 for other models). As we can see from Figure 5, the accuracy of the contact prediction yields a modest correlation (PCC=0.23) with 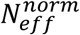 (log scale), in which targets with higher 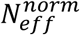 tend to have higher contact prediction accuracies. However, for about 60% (13/22) of the targets with very low 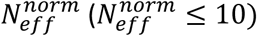, the top L/5 long-range contacts predicted by DRN-1D2D still have an accuracy higher than 50%.

We further grouped the targets into two groups according to the value of 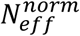 (low 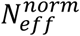 group: 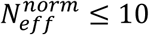; high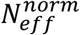 group: 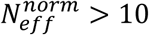), and analyzed the performances of DRN-1D2D and other models on the two groups of targets respectively. In Figure 6, we show the mean precisions of the top L/5 and top L/2 of the medium + long-range (Figure 6A) and long-range (Figure 6B) contacts predicted by each model on the two groups of targets. As we can see from Figure 6, for both the two groups, DRN-1D2D consistently achieved the best performances among these models. Besides, if BCE-2D is considered as our baseline model (i.e. the model without using the dimensional hybrid residual block and the singularity enhanced loss), the performance improvement of DRN-1D2D on the low 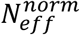 group is much higher than that on the high 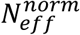, which illustrates that DRN-1D2D has a lower dependency on the number of sequences in the MSA (also see Table S2).

### 3.4 The performance of DRN-1D2D on large protein contact prediction and extra long-range contact prediction

In the development of DRN-1D2D, a dimensional hybrid residual block was introduced to increase the effective receptive field of the network, and a new loss function was proposed to emphasize the large proteins and the contacts in the contact-sparser regions of the contact maps. Therefore, it is expected that the contact prediction for large proteins and contact-sparser regions can be enhanced through these changes. To validate this assumption, we analyzed the contact prediction performance of DRN-1D2D on large proteins and extra long-range regions (sequence distance ≥ 50).

We re-grouped the CASP targets into two groups according to their sequence lengths (small protein group: L ≤ 150; large protein group: L > 150). In Figure 7, we show the mean precisions of the top L/5 and top L/2 of the medium + long-range (Figure 7A) and long-range (Figure 7B) contacts predicted by each model on the two groups of targets. As we can see from Figure 7, for all the models, the contact prediction precisions on the large proteins tend to be higher than those on the small proteins (also see Figure S4), which is consistent with the CASP observations [2]. However, on both the two groups, DRN-1D2D consistently achieved the best performance. Besides, when comparing with our baseline model BCE-2D, the performance improvement on the large proteins is much higher than that on the small proteins (also see Table S3), which supports our assumption that the two innovations in the model development can enhance the contact prediction for large proteins.

We further analyzed the performance of DRN-1D2D on extra long-range contact prediction. The contacts in extra long-range regions of protein contact maps are generally very sparse, therefore, predicting extra long-range contacts is quite challenging. Actually, we noticed that four targets of CASP13 (T0957S2-D1,T0960-D2,T0963-D2,T0991-D1) were removed from the CASP official evaluation of the extra long-range contact prediction for they have very few number of extra long-range contacts (the numbers of the extra long-range contacts of the four targets are 2, 11, 9, 2 respectively). Thus, these four targets were excluded from our analysis as well. In Table S3, we show the mean precisions of the extra long-range contacts predicted by different models. As we can see from the table, DRN-1D2D achieved best performance in most of the cases, with the only exception being the top L/5 predictions of CASP13 dataset, in which the precision of DRN-1D2D is slightly lower than TripleRes (1%) and RaptorX-Contact (2%). Comparing with the medium + long-range and long-range contact predictions, the gaps of the mean precisions between DRN-1D2D and other top models are quite small (1%-2%). This is mainly caused by the existence of several extremely challenging targets, for which almost all the models totally failed in the extra long-range contact prediction (see Data S1). However, if we compare the performance of DRN-1D2D with the baseline model BCE-2D, the improvement is still dramatic (2%-6%) and systematic (see Figure S5). Therefore, the result still supports that the extra long-range contact prediction can be enhanced through the two innovations in the model development.

## 4. Conclusion

A new deep residual learning-based protein contact prediction model referred to as DRN-1D2D was presented in this study. Different from previous models, a new type of residual block hybridizing 1D and 2D convolutions was applied to build the residual network of DRN-1D2D, with which we show the effective receptive field of the residual network can be significantly increased, and a new loss function emphasizing the easily misclassified residue pairs was applied to enhance the model training. We first evaluated DRN-1D2D on Xu’s original three test sets together with two in house-developed reference models: one based on the basic 2D residual block and the BCE loss function, and another based on the dimensional hybrid residual block and the BCE loss function. Besides, Xu’s original model performance was also used as a reference. The result shows that DRN-1D2D consistently outperforms the two reference models and Xu’s original model. Between the two reference models, the model based on the new residual block consistently outperforms the basic 2D residual block, which confirms that the introduction of the dimensional hybrid residual block and the singularity enhanced loss function are effective protocols to improve the model performance for protein contact prediction. We further evaluated DRN-1D2D on recent CASP targets together with the two reference models and six state-of-the-art protein contact prediction models including DeepCov, DeepCon, DeepConPred2, SPOT-contact, RaptorX-Contact and TripleRes. The result shows that DRN-1D2D consistently achieved the best performance among these models, although DRN-1D2D was trained on a relative older and smaller dataset and has a much smaller network architecture than RaptorX-Contact and TripleRes. It is reasonable to assume our model can be further improved if we can build a deeper residual network with more residual blocks and train the model on larger training sets.

## Key points

- A dimensional hybrid residual block is designed to improve the effective receptive field of the residual network.
- A singularity enhanced loss function is proposed to enhance the model training.
- Both the dimensional hybrid residual block and the singularity enhanced loss function can be employed to improve the model performance for protein contact prediction.
- The developed model consistently and significantly outperforms the state-of-the-art protein contact prediction models.

## Availability

DRN-1D2D is available at https://github.com/ChengfeiYan/DRN-1D2D.

## Funding

This work is supported by the startup grant of Huazhong University of Science and Technology.

**Yunda Si** is a PhD student in the School of Physics at Huazhong University of Science and Technology. His research interests include protein structure prediction, protein-protein interaction prediction and deep learning.

**Chengfei Yan** is an associate professor in the School of Physics at Huazhong University of Science and Technology. His research interests include molecular docking, protein-protein interaction prediction and biological data mining.

