## Supplemental Figures and Tables for "Improved protein contact prediction using dimensional hybrid residual networks and singularity enhanced loss function"

**Supplementary**

Text S1: Preparing the input features of DRN-1D2D

For each target in the CASP13 and CASP14 datasets, we generated the input features of DRN-1D2D including the position-specific scoring matrix (PSSM), the predicted 2-state secondary structure matric (SS3), the predicted 3-state solvent accessibility matrix (ACC), the evolutionary coupling matrix, the mutual information matrix, the APC-corrected mutual information matrix and the pairwise contact potential matrix from the MSA of its homologous proteins according to the protocols described by Xu ( (https://github.com/j3xugit/RaptorX-Contact). The technical details of the protocols are also described as follows：

(1) PSSM. We first used HHMake from HH-suite[1] to build an HHM profile from the provided MSA, and then we used the script LoadHMM.py from RaptorX-contact[2] to generate the PSSM from the HMM profile.
(2) ACC &SS3. We used RaptorX-Property[3] to predict ACC and SS3 from the provided MSA.
(3) Evolutionary coupling matrix. We ran CCMpred[4] with the option "-R" on the provided MSA to generate the evolutionary coupling matrix. "-R" was set to normalize the output matrix by its mean and standard deviation.
(4) The mutual information matrix, the APC-corrected mutual information matrix and the pairwise contact potential. We applied alnstats from MetaPSICOV[5] to generate the three pairwise matrices.

Table S1. Parameter number and ERF comparison for residual networks formed by different numbers of 2D blocks and 1D2D blocks

| Block | 2D Block | | | 1D2D Block |
|---|---|---|---|---|
| # of blocks | 25 | 75 | 142 | 25 |
| Params | 1 | 3 | 5.68 | 3 |
| ERF | 1 | 1.76 | 2.435 | 2.426 |

Note: The residual network formed by 25 2D blocks are used a reference. The ERF size of each residual network is evaluated by the full width at half maxima (FWHM) of the impact distribution. For example, the ERF of the residual network formed by 25 1D2D blocks has an FWHM 2.426 times larger than the residual network formed by the same number of 2D blocks.

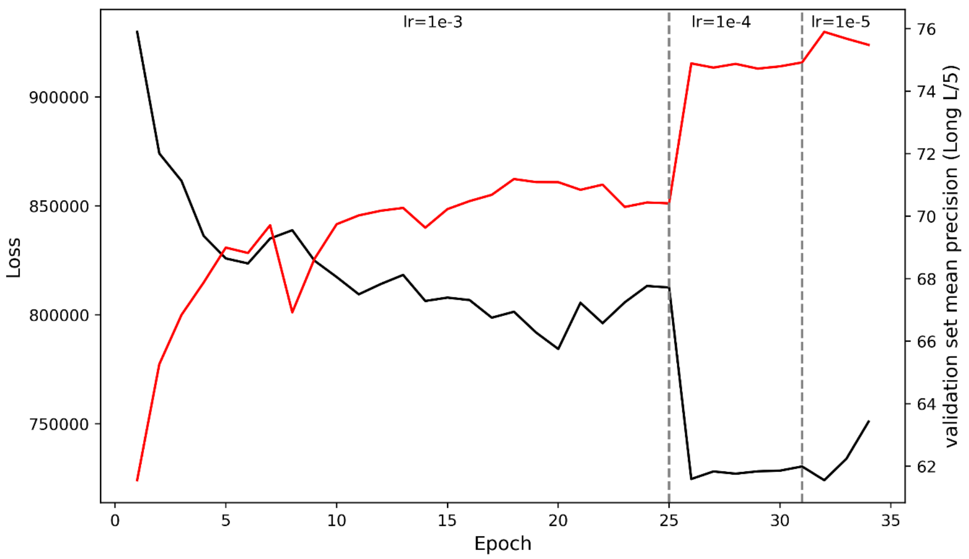


Figure S1. The change of training loss, validation accuracy and learning rate for a model training of DRN-1D2D.

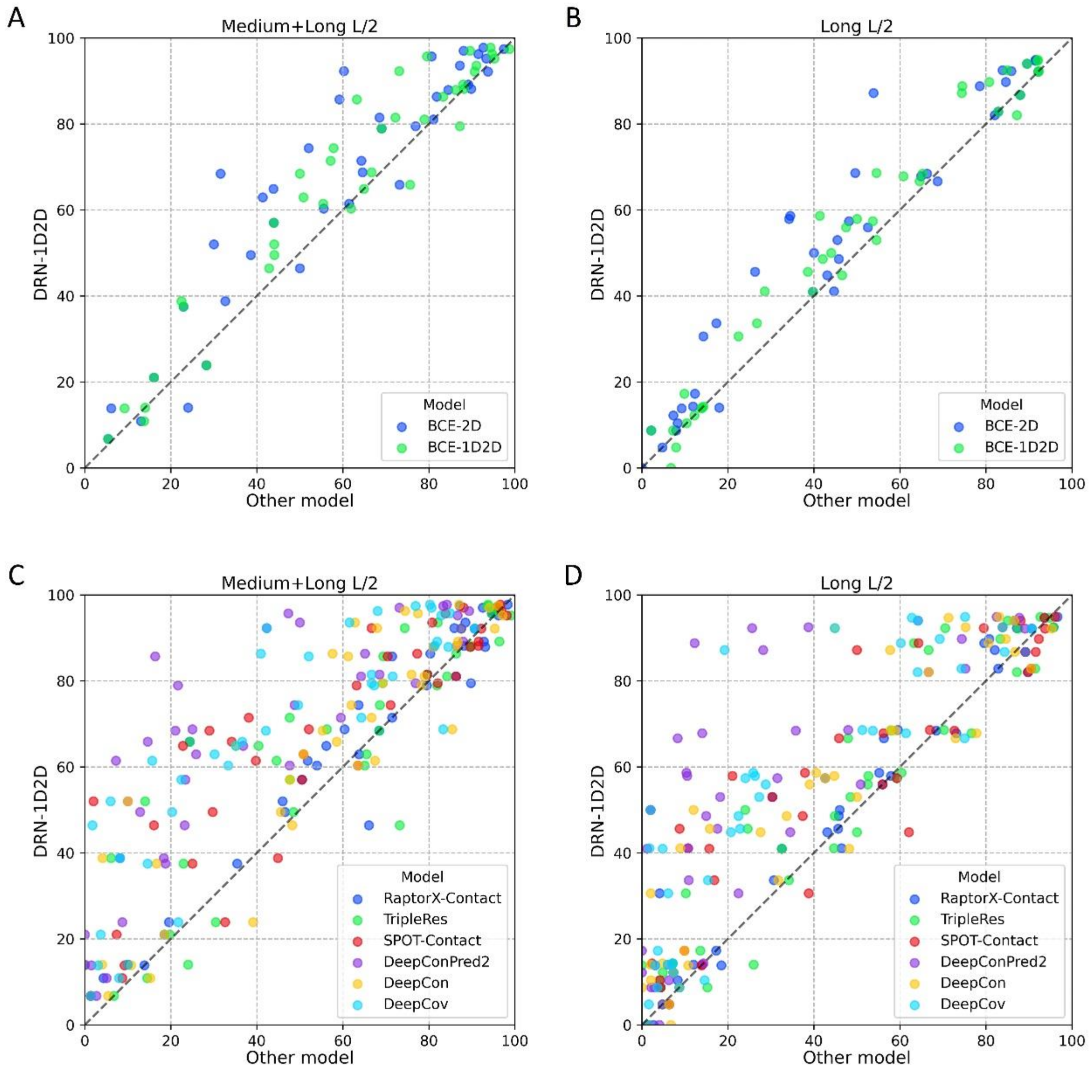


Figure S2. The performance comparison between DRN-1D2D and other models for all individual targets from the CASP datasets. (A) DRN-1D2D versus the two in house-developed reference models for top L/2 medium + long-range predicted contacts. (B) DRN-1D2D versus the two in house-developed reference models for top L/2 long-range predicted contacts (C) DRN-1D2D versus six state-of-the-art outside models for top L/2 medium + long-range predicted contacts. (D) DRN-1D2D versus six state-of-the-art outside models for top L/2 long-range predicted contacts.

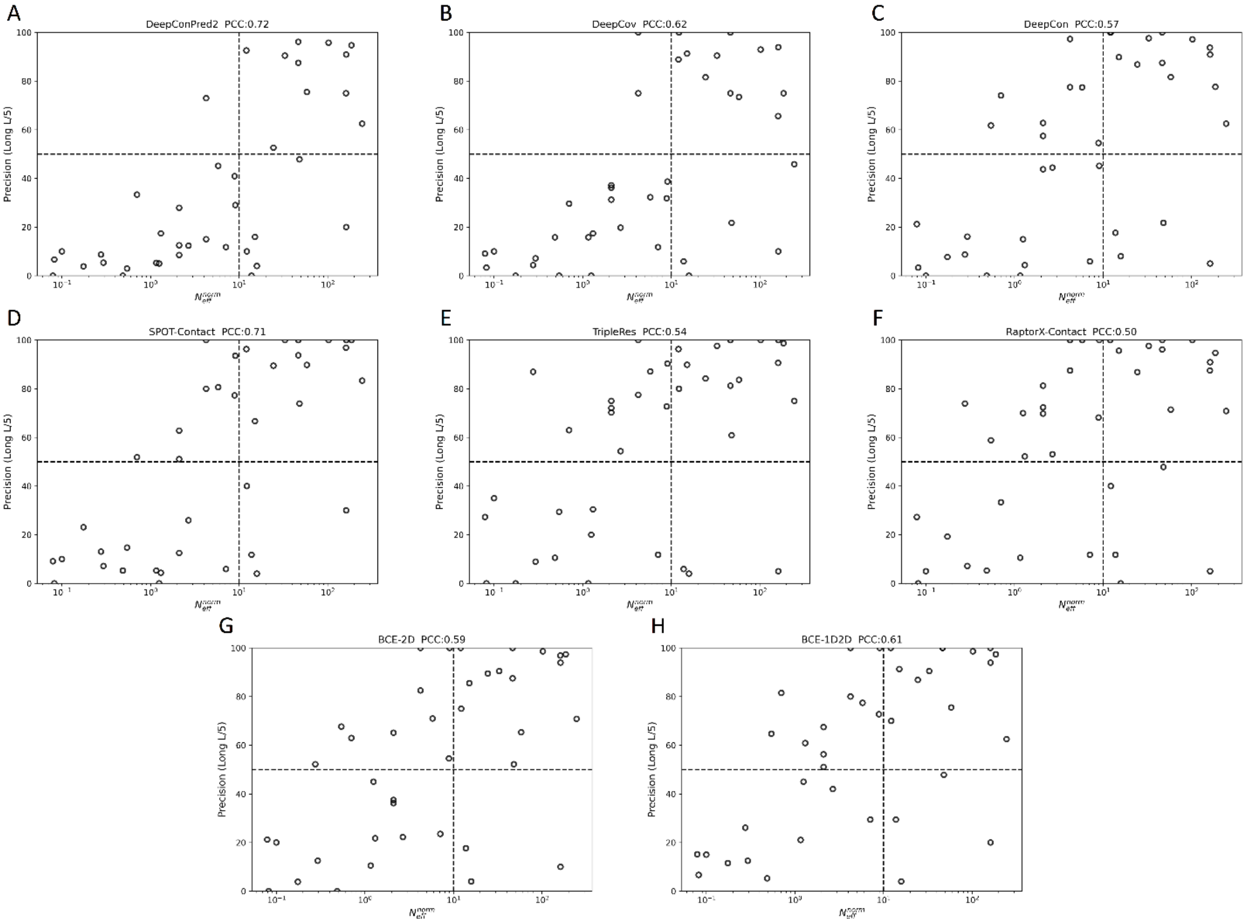


Figure S3. Precision of top L/5 long-range predicted contacts versus $N_{eff}^{norm}$ of MSAs for different models: (A) DeepConPred2; (B) DeepCov; (C) DeepCon; (D) SPOT-Contact; (E) TripleRes; (F) RaptorX-Contact; (G) BCE-2D; (H) BCE-1D2D.

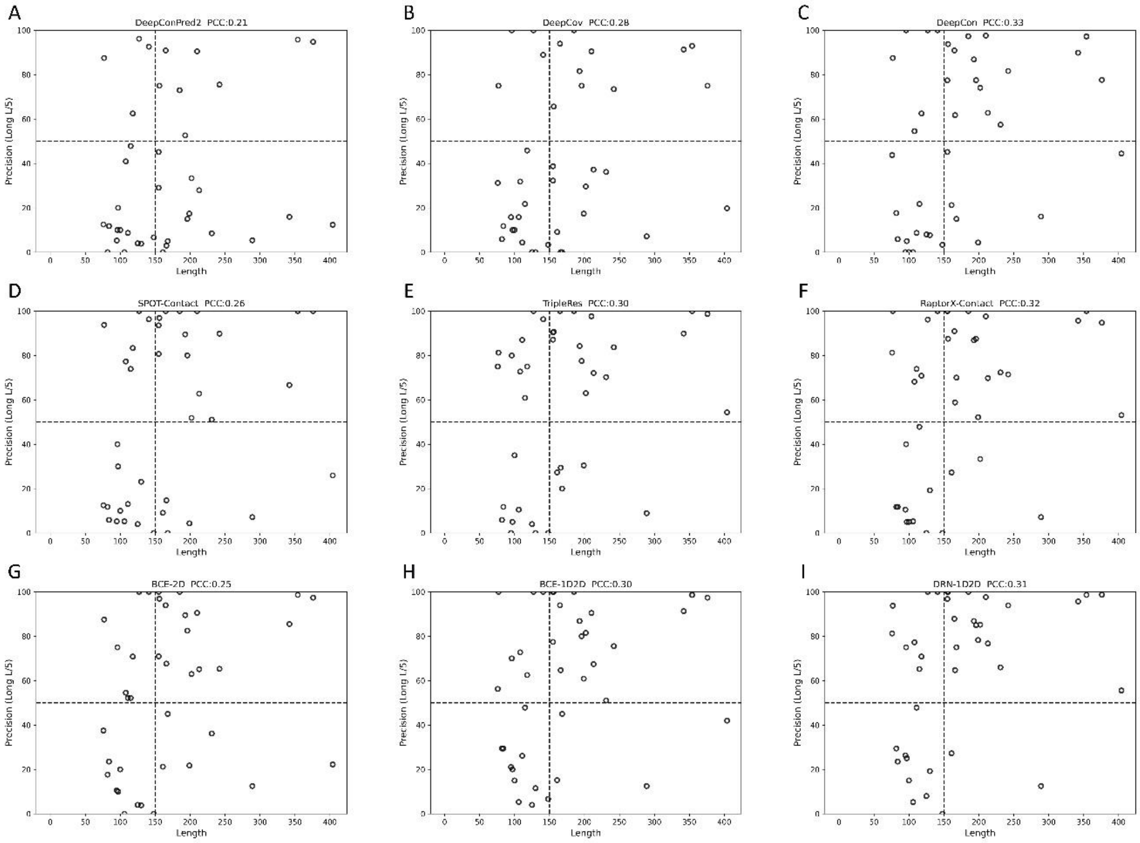


Figure S4. Precision of top L/5 long-range predicted contacts versus protein sequence length for different models: (A) DeepConPred2; (B) DeepCov; (C) DeepCon; (D) Spot-Contact; (E) TripleRes; (F) RaptorX-Contact; (G) BCE-2D; (H) BCE-1D2D; (I) DRN-1D2D.

**Table S2. The improvements of DRN-1D2D over the baseline model BCE-2D on the two groups of targets with $N_{eff}^{norm} \leq 10$ and $N_{eff}^{norm} > 10$ respectively**

| Model | 0-10 | | | | 10+ | | | |
|---|---|---|---|---|---|---|---|---|
| | Medium+Long | | Long | | Medium+Long | | Long | |
| | L/5 | L/2 | L/5 | L/2 | L/5 | L/2 | L/5 | L/2 |
| BCE-2D | 0.52 | 0.41 | 0.39 | 0.32 | 0.89 | 0.78 | 0.74 | 0.64 |
| BCE-1D2D | 0.55 | 0.44 | 0.45 | 0.35 | 0.89 | 0.80 | 0.76 | 0.65 |
| DRN-1D2D | 0.64 | 0.53 | 0.53 | 0.40 | 0.91 | 0.82 | 0.79 | 0.68 |
| Δ | 0.12 | 0.12 | 0.14 | 0.08 | 0.03 | 0.04 | 0.05 | 0.04 |

**Table S3. The improvements of DRN-1D2D over the baseline model BCE-2D on the two groups of targets with sequence length $\leq 150$ and sequence length $> 150$ respectively**

| Model | Target length: 0-150 | | | | Target length: 150+ | | | |
|---|---|---|---|---|---|---|---|---|
| | Medium+Long | | Long | | Medium+Long | | Long | |
| | L/5 | L/2 | L/5 | L/2 | L/5 | L/2 | L/5 | L/2 |
| BCE-2D | 0.63 | 0.53 | 0.40 | 0.35 | 0.74 | 0.63 | 0.68 | 0.57 |
| BCE-1D2D | 0.62 | 0.54 | 0.43 | 0.37 | 0.78 | 0.67 | 0.73 | 0.59 |
| DRN-1D2D | 0.67 | 0.58 | 0.48 | 0.39 | 0.84 | 0.74 | 0.80 | 0.65 |
| Δ | 0.04 | 0.05 | 0.08 | 0.04 | 0.10 | 0.11 | 0.12 | 0.08 |

**Table S4. The mean precisions of extra long-range (sequence distance $\geq 50$) contacts predicted by different models on CASP 13 and 14 targets**

| Method | Extra-Long | | | | | |
|---|---|---|---|---|---|---|
| | CASP13 | | | CASP14 | | |
| | L/5 | L/2 | L | L/5 | L/2 | L |
| DeepCov | 0.53 | 0.40 | 0.28 | 0.19 | 0.15 | 0.11 |
| DeepCon | 0.62 | 0.47 | 0.33 | 0.29 | 0.21 | 0.16 |
| DeepConPred2 | 0.39 | 0.32 | 0.24 | 0.16 | 0.11 | 0.08 |
| SPOT-Contact | 0.67 | 0.52 | 0.36 | 0.22 | 0.16 | 0.12 |
| BCE-2D | 0.67 | 0.50 | 0.36 | 0.27 | 0.20 | 0.15 |
| TripleRes | **0.71** | 0.55 | 0.39 | 0.28 | 0.21 | 0.16 |
| BCE-1D2D | 0.65 | 0.52 | 0.37 | 0.27 | 0.21 | 0.15 |
| RaptorX-Contact | 0.70 | 0.55 | 0.39 | 0.30 | 0.23 | 0.16 |
| DRN-1D2D | 0.69 | **0.56** | **0.40** | **0.31** | **0.24** | **0.18** |

Note: 1. T0957S2-D1, T0960-D2, T0963-D2, T0991-D1 from CASP13 were excluded from the evaluation of the extra long-range contact prediction for they have very few number (2, 11, 9, 2) of extra long-range contacts. These four targets were also excluded in the CASP official evaluation of the extra long-range contact prediction.

2. The highest accuracy in each category is highlighted in bold font.

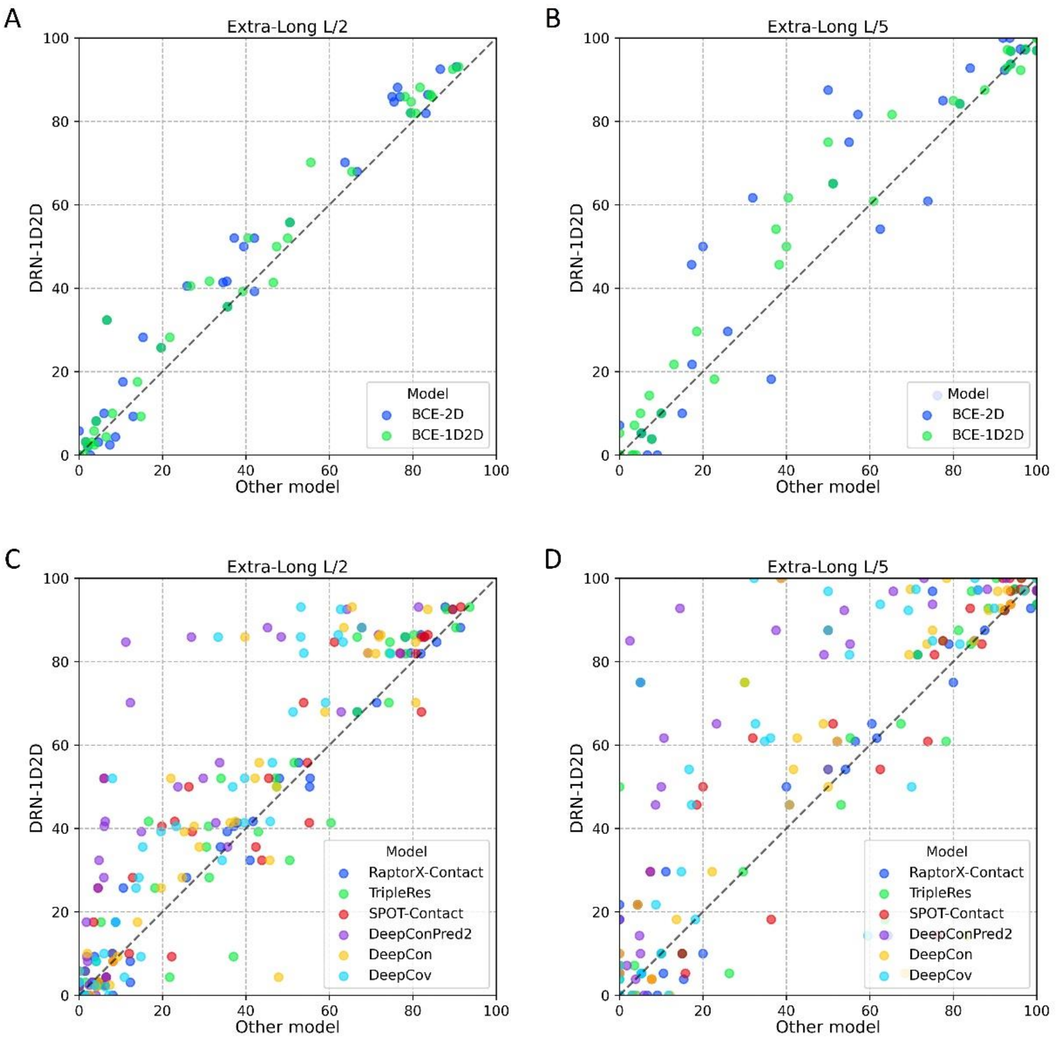


Figure S5. The performance comparison on extra long-range contact prediction between DRN-1D2D and other models for all individual targets of the CASP 13 and 14 free modeling targets. (A) DRN-1D2D versus the two in house-developed reference models for top L/2 extra long-range predicted contacts. (B) DRN-1D2D versus the two in house-developed reference models for top L/5 extra long-range predicted contacts (C) DRN-1D2D versus six state-of-the-art outside models for top L/2 extra long-range predicted contacts. (D) DRN-1D2D versus six state-of-the-art outside models for top L/5 extra long-range predicted contacts.

**Data S1. The detailed performances of different protein contact prediction models on each individual target from the CASP 13 and CASP 14 datasets.**
